# Sex-focused analyses of M83 A53T hemizygous mouse model with recombinant human alpha-synuclein preformed fibril injection identifies female resilience to disease progression: A combined magnetic resonance imaging and behavioural study

**DOI:** 10.1101/2024.05.24.595642

**Authors:** Stephanie Tullo, Janice Park, Daniel Gallino, Megan Park, Kristie Mar, Vladislav Novikov, Rodrigo Sandoval Contreras, Raihaan Patel, Esther del Cid-Pellitero, Edward A. Fon, Wen Luo, Irina Shlaifer, Thomas M. Durcan, Marco A.M. Prado, Vania F. Prado, Gabriel A. Devenyi, M. Mallar Chakravarty

## Abstract

Alpha-synuclein (aSyn) pathology has been extensively studied in mouse models harbouring human mutations. In spite of the known sex differences in age of onset, prevalence and disease presentation in human synucleinopathies, the impact of sex on aSyn propagation has received very little attention. To address this need, we examined sex differences in whole brain signatures of neurodegeneration due to aSyn toxicity in the M83 mouse model using longitudinal magnetic resonance imaging (MRI; T1-weighted; 100 μm^3^ isotropic voxel; acquired −7, 30, 90 and 120 days post-injection [dpi]; n≥8 mice/group/sex/time point). To initiate aSyn spreading, M83 mice were inoculated with recombinant human aSyn preformed fibrils (Hu-PFF) or phosphate buffered saline (PBS) injected in the right dorsal striatum. We observed more aggressive neurodegenerative profiles over time for male M83 Hu-PFF-injected mice when examining voxel-wise trajectories. However, at 90 dpi, we observed widespread patterns of neurodegeneration in the female Hu-PFF-injected mice. These differences were not accompanied with any differences in motor symptom onset between the male and female Hu-PFF-injected mice. However, male Hu-PFF-injected mice reached their humane endpoint sooner. These findings suggest that post-motor symptom onset, even though more accelerated disease trajectories were observed for male Hu-PFF-injected mice, neurodegeneration may appear sooner in female Hu-PFF-injected mice (prior to motor symptomatology). These findings suggest that sex-specific synucleinopathy phenotypes urgently need to be considered to improve our understanding of neuroprotective and neurodegenerative mechanisms.

## 1. Introduction

Synucleinopathies, including Parkinson’s disease (PD), Dementia with Lewy Bodies (DLB), Multiple systems atrophy (MSA), pure autonomic failure and REM sleep behavior disorder (RBD), are neurodegenerative disorders that share a common feature of aberrant accumulation of alpha-synuclein (aSyn) aggregates in the brain (Boeve et al., 2003; Jellinger, 2003; Kao et al., 2009), which may be underlying the mechanisms of pathogenesis via aSyn spreading (Polymeropoulos et al., 1997; Spillantini et al., 1997; Calabresi et al., 2023). To this end, evidence of cell-to-cell propagation has supported the hypothesized prion-like spreading of aSyn that contributes to neurotoxicity and downstream neurodegeneration. Evidence for the prion-like spreading of aSyn emerged from post-mortem analysis of patients’ brains that received fetal dopaminergic cell transplantation in the substantia nigra pars compacta (Kordower et al., 2008; Li et al., 2008) showing host-to-graft aSyn transfer. Subsequent cellular assays of aSyn cell-to-cell transfer (Bétemps et al., 2014; Desplats et al., 2009; Fares et al., 2016; Panattoni et al., 2018; Tapias et al., 2017; Volpicelli-Daley et al., 2014) and animal models of aSyn spreading (Froula et al., 2019; Lackie et al., 2022; Luk et al., 2012a; Luk et al., 2012b; Masuda-Suzukake et al., 2013; Masuda-Suzukake et al., 2014; Mougenot et al., 2012; Sacino et al., 2014; Sorrentino et al., 2022; Tullo et al., 2023; Watts et al., 2013) further support the prion-like hypothesis.

Higher prevalence, incidence, increased disease severity and susceptibility in men has been reported in PD (Picillo et al., 2017), MSA (Coon et al., 2019; Yamamoto et al., 2014) and DLB (Utsumi et al., 2020), and even in PD prodromes, like RBD (Zhou et al., 2015). The compelling clinical evidence clearly underscores the need to examine the role of sex in disease progression and presentation (Choleris et al., 2018; de Lange et al., 2021; Eid et al., 2019; Raheel et al., 2023). Yet sex differences in the prion-like spreading of aSyn and the potential downstream consequences are not clearly understood. This is not altogether surprising as sex is often neglected as a biological variable in preclinical research (Galea et al., 2020; Galea & Parekh 2023; Galea et al., 2023). This limitation extends to using the M83 mouse model, harbouring an A53T mutation in the human aSyn transgene, that is commonly used to examine the prion-like spreading hypothesis (Bétemps et al., 2014; Froula et al., 2019; Lackie et al., 2022; Luk et al., 2012a; Luk et al., 2012b; Masuda-Suzukake et al., 2013; Masuda-Suzukake et al., 2014; Mougenot et al., 2012; Sacino et al., 2014; Sorrentino et al., 2022; Tullo et al., 2023; Watts et al., 2013). Aside from a recent study that used intramuscular injection (Sorrentino et al., 2022) and a magnetic resonance imaging (MRI) study from our group (Tullo et al., 2023), few others using the M83 mouse line have explicitly explored sex differences. Given the cross-sectional nature of our recent work, it is likely that a longitudinal design may be more sensitive to inter-individual variation that results in sex differences (Lerch et al. 2012; Valiquette et al. 2023). Regardless of sex, most studies using M83 mice characterize aSyn inclusions at a single time point to characterize aSyn spread. Taken together, there is an urgent need to understand sex-specific trajectories using longitudinal study designs to better characterize the sex-specific prion-like spreading of aSyn to better model PD-like pathophysiology.

To this end, we studied the M83 transgenic mice injected with human aSyn preformed fibrils (Hu-PFF) in the right dorsal striatum. We examined sex-specfic trajectories of the disease time course using longitudinal structural MRI (Borg & Chereul, 2008; Gallino et al., 2019; Guma et al., 2021; Johnson et al., 2007; Kong et al., 2018; Lerch et al., 2011; Pagani et al., 2016; Rollins et al., 2019). Next, we derived sex-specific whole brain atrophy patterns at specific focal time points, pre-/peri-motor symptom onset and post-motor symptom onset where the mice neared their humane endpoint. Our study significantly contributes to the expanding body of knowledge, offering insights that could profoundly impact the design and implementation of sex-specific therapeutic interventions for synucleinopathies.

## 2. Materials and Methods

### 2.1. Experimental Design

The experimental timeline is shown in Figure 1 and group numbers in Supplementary Table S1. We examined groups receiving Hu-PFF or PBS in the right dorsal striatum at 7 days prior to the injection (−7 dpi) for a baseline assessment, 30 dpi (earliest signs of aSyn Lewy-body like pathology (Luk et al., 2012)), 90 dpi (when motor symptomatology onset is typically observed (∼ 90-100 dpi) (Bétemps et al., 2014; Luk et al., 2012a; Sacino et al., 2014; Sorrentino et al., 2022; Tullo et al., 2023; Watts et al., 2013)), and 120 dpi (when the mice are fully symptomatic and nearing the end stage of the disease. Mice were studied at specified time points ± 2 days. Prior to data collection, all mice were weighed. Mice underwent *in vivo* MRI followed by the administration of behavioural tasks on subsequent days. Motor behaviours were assayed with pole test (Matsuura et al., 1997), rotarod (Janickova et al., 2017), and wire hang (Froula et al. 2019) to assess motor deficits over time. All experiments started with approximately 30 mice per group at −7 days post-injection, with decreasing numbers at each subsequent time point, as mice were diverted at each time point for future terminal experimental methods with a final sample of ∼8 mice/sex/group at 120 dpi. We examined sex differences in voxel-wise trajectories of neurodegeneration, neurodegenerative patterns at the two last time points, and phenotypic differences in M83 mice after receiving Hu-PFF inoculation. Finally, we examined whether there were inherent sex differences in the M83 hemizygous mice (not injected), specifically with regards to differences in their normative aSyn expression, as determined by western blotting.

**Figure 1.**
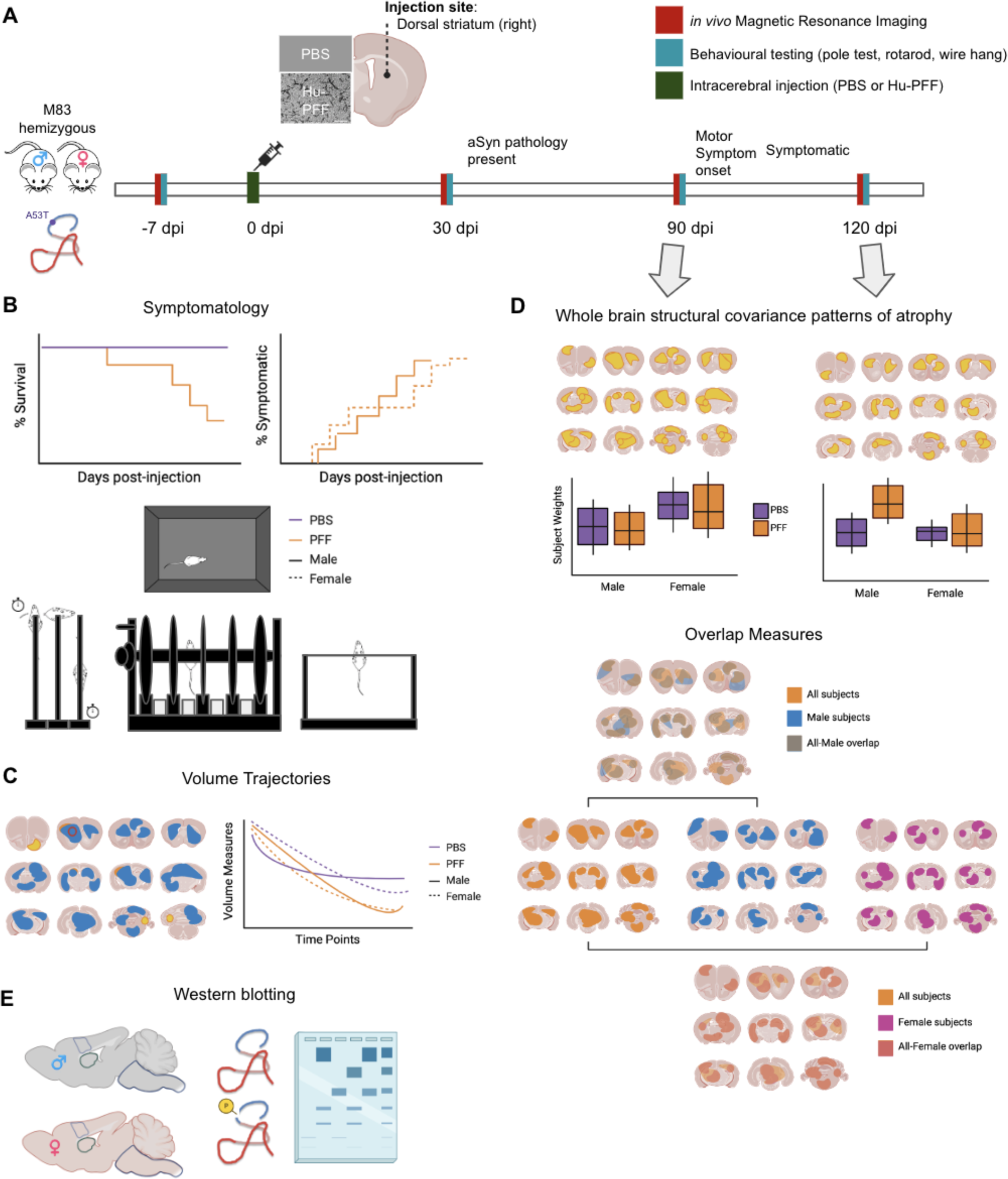
Experimental overview with conceptual figures (not displaying real data). [A] Male and female hemizygous M83 mice were injected with either PBS or Hu-PFF; approximately 8 mice per injection group per sex per time point with mice siphoned at each time point. The mice underwent in vivo MRI and motor behavioural testing (pole test, rotarod, and wire hang test) at four time points: −7, 30, 90 (peri-motor symptom onset), and 120 (post-motor symptom onset) days post-injection (dpi). [B] Motor symptomatology was assessed via visual observation of the mice ambulating in their home cage, as well as via their performance on the pole test, rotarod and wire hang test. [C] Univariate analyses were performed to assess group differences in neuroanatomical trajectories across the number of days post-injection. [D] Subsequently, multivariate analysis (orthogonal projective non-negative matrix factorization) was performed to examine whole brain aSyn Hu-PFF-induced atrophy patterns using voxel-wise measures of volume (Jacobians derived from deformation-based morphometry) at 90 and 120 dpi. Sex differences were evaluated by either examining the injection group by sex differences in NMF subject weights or assessing the overlap between spatial patterning for each sex-specific pattern and the all subjects pattern. [E] Western blotting experiments were performed on non-injected M83 hemizygous mice to examine normative sex differences in these transgenic mice with regards to aSyn expression.

### 2.2. Animals

We used transgenic hemizygous M83 mice (B6; C3H-Tg[SNCA]83Vle/J) bred in-house (F4-6), expressing one copy of human alpha-synuclein bearing the familial PD-related A53T mutation under the control of the mouse prion protein promoter (TgM83+/-) (Giasson et al., 2002), in addition to the endogenous mouse alpha-synuclein, maintained on a C57BL/C3H background.The M83 hemizygous mice used here typically present no motor phenotype until 22 to 28 months of age (Giasson et al., 2002). aSyn Hu-PFFs, synthesized and characterized by the Early Drug Discovery Unit (EDDU) at the Montreal Neurological Institute (production: https://zenodo.org/record/3738335#.Yp46fXbMKUl; characterization: https://zenodo.org/record/3738340#.YzIjBezML0p), were injected in the right dorsal striatum to trigger accumulation of toxic aSyn and accelerate symptom onset (Luk et al., 2012a). All study procedures were performed in accordance with the Canadian Council on Animal Care and approved by the McGill University Animal Care Committee, and the University of Western Ontario (2020-162; 2020-163). For more information, see section *7.1. Animals* in the Supplementary Materials.

### 2.3 Injection material and Stereotaxic injections

Healthy 3 to 4-month-old hemizygous M83 mice were injected with Hu-PFF or phosphate-buffered saline (PBS) in the right dorsal striatum, consistent with previous work modelling known disease epicentres of spreading (Bétemps et al., 2014; Luk et al., 2012a; Mougenot et al., 2012; Tullo et al., 2023). Groups were randomly assigned, with an equal number of mice across injection assignment and sex. Post-injection, experimenters were blind to the injection assignment of each mouse during the data acquisition processes and during statistical analysis. For more information on *Injection material and Stereotaxic injections,* see Supplementary Materials, section *7.2. Injection materials and Stereotaxic injections*.

### 2.4 Magnetic Resonance Imaging (MRI) acquisition

Mice underwent MRI at −7, 30, 90 and 120 dpi, unless they reached a humane endpoint prior. MRI acquisition was performed on 7.0-T Bruker Biospec (70/30 USR; 30-cm inner bore diameter; AVANCE electronics) at the Cerebral Imaging Centre of the Douglas Research Centre (Montreal, QC, Canada). In vivo T1-weighted images (FLASH; Fast Low Angle SHot) were acquired for each subject at each time point (TE/TR of 4.5 ms/20 ms, 100 μm isotropic voxels, 2 averages, scan time= 14 min, flip angle=20°) (see section *7.3. MRI acquisition: anaesthesia protocol* in the Supplementary Materials for more details).

#### 2.4.1 Image processing: Deformation-based morphometry

After scanning all images underwent pre-processing and quality control (See Supplementary Materials, section *7.4. Pre-processing*) prior to being used as input in the deformation morphometry pipeline. Here, subjects are aligned with a series of linear and nonlinear registration steps with iterative template refinement (Avants et al. 2011) to generate a study-specific template enabling voxel correspondence across every subject timepoint (antsMultivariateTemplateConstruction2.sh; https://github.com/CoBrALab/twolevel_ants_dbm). This method has been previously used by our group to examine voxel-wise volumetric change over time (Gallino et al., 2019; Guma et al., 2021; Kong et al., 2018; Rollins et al., 2019) (see section *7.5. Deformation-based morphometry* in the Supplementary Materials for more details). The outputs were inspected to assess quality control of the registration by visually assessing the resultant images of the registrations for a proper orientation, size, and sensible group average, prior to performing the analyses (https://github.com/CoBrALab/documentation/wiki/Mouse-QC-Manual-(Structural)).

### 2.5. Motor behaviour tasks

After scanning at each of the four MRI timepoints (−7, 30, 90, 120 dpi), the mice underwent a behavioural test each day for three days (>24 hours post-scan) to quantitatively measure motor symptomatology; namely: pole test, rotarod and wire hang test. Tests were conducted during the light phase, between 8 am and 8 pm. The mice were habituated to the testing room for one hour prior to administering the behavioural test (see Supplementary Materials, section *7.6. Behavioural Tasks* for more details).

### 2.6. Western blotting

Cortical, striatal, and brainstem tissue from 6-7 months old non-injected M83 hemizygous mice was homogenized to obtain radioimmunoprecipitation assay (RIPA) soluble and insoluble fractions to examine differences in aSyn expression between the sexes. The following primary antibodies were used: phospho S129 (1:1000, Cat# ab51253, Abcam, RRID:AB_869973), anti-human a-Syn (1:1000, Cat# ab27766, Abcam, RRID:AB_727020), anti-alpha synuclein (1:1000, Cat# 610787, BD Biosciences, RRID: AB_398108), anti-actin HRP (1:25,000, Cat#A3854, Sigma-Aldrich, RRID:AB_262011). Densitometry was processed and analyzed using Bio-Rad ImageLab software. For more information see the Supplementary Materials, section *7.7. Western Blotting*.

### 2.7. Statistical analyses

#### 2.7.1. Disease progression and motor symptomatology analysis

Disease progression (in terms of survival) and motor symptom onset were examined using Cox Proportional Hazard modeling (Kassambara et al., 2018) (see Supplementary Materials; *7.8. Cox Proportional Hazard modelling*). We also examined differences in average dpi overt motor symptomatology was first observed in ambulating mice using general linear model (average dpi ∼ injection_group*sex). For longitudinal measures such as weight and rotarod performance (averaged across trials at each time point), linear mixed effect models were used to assess injection group by sex differences across the time points (∼ injection_group*sex*time point + (1|subject_ID)). For pole test and wire hang tests, performance was examined with regards to latency and failure/success rate using a Cox Proportional Hazard model (Kassambara et al., 2018) as these two tasks have strict time cut-offs to determine the success/fail rate. Bonferroni correction was performed to account for multiple comparisons (p=0.05/8; 2 motor tests and 4 time points). Statistical analyses were carried out using R software (3.5.0) with the RMINC package (https://github.com/Mouse-Imaging-Centre/RMINC) for voxel-wise analysis, and the survival package (https://cran.r-project.org/web/packages/survival/index.html) for the cox modelling.

#### 2.7.2. Longitudinal volumetric analysis

We performed univariate analyses to assess group-level voxel-wise differences in neuroanatomical change using the relative Jacobian determinants (exclusively modelling the nonlinear transformations, representing volume differences, that were log transformed and blurred using Gaussian smoothing) generated from DBM using linear mixed effects models at each voxel to examine volume differences between the injection groups and sex of the mice over time:

Relative volume at each voxel ∼ injection_group*sex*time point + (1| subject_ID) Statistical findings were corrected with False Discovery Rate (FDR) (Benjamini and Hochberg 1995). Statistical analyses were carried out using R software (3.5.0) and the RMINC package (https://wiki.phenogenomics.ca/display/MICePub/RMINC).

#### 2.7.3. Whole brain structural covariance analysis

##### 2.7.3.1 Orthogonal projective non-negative matrix factorization

We previously demonstrated that modelling MRI-derived atrophy in M83 mice with striatal inoculation using methods that capture spatial covariance patterns (Tullo et al., 2023) resulted in substantial homology to human patients with PD (Yau et al., 2018; Zeighami et al. 2015; 2019). In our previous work we used a multivariate technique known as orthogonal projective non-negative matrix factorization (OPNMF) (Patel et al., 2020; Robert et al., 2022; Sotiras et al., 2015; Yang & Oja, 2010).

We used OPNMF to decompose an input matrix of z-score normalized inverted absolute Jacobian determinants for each voxel into two output matrices. The first is component weights describing the relative contribution of each voxel to a covariance pattern that can be visualized on the mouse brain template to characterize neuroanatomical localization. The second contains subject-specific weights for each component that can be used to understand inter-individual differences (see section *7.9. Orthogonal projective non-negative matrix factorization* in Supplementary Materials). Importantly, the methodology contains no information (treatment nor sex) regarding the mice under investigation. General linear models were used to examine the sex by injection group interaction in the subject weights post-hoc for each component to identify sex differences.

##### 2.7.3.1.2. Sex differences in structural covariance analysis

Typically, examination of the OPNMF subject weights has been used to examine inter-individual variation. However, since the patterns that are elucidated are threshold free and are sparse matrices, they can be used to evaluate the overlap between the pattern that best differentiates the Hu-PFF- and saline-injected groups as well as the male- and female-specific aSyn-mediated neurodegeneration to better understand sex-specific overlap with the group pattern. We measured the overlap between sex-specific and whole group (all subjects) binarized maps to assess pattern shape and extent using the Dice similarity coefficient (also referred to as Dice’s Kappa) (κ) metric. This is similar to recent work from our group used to quantify network shape in rodent resting state functional MRI (Desrosiers-Gregoire et al., 2024).

The Dice’s Kappa overlap metric score is: κ=2a/(2a+b+c); where a is the number of voxels common to comparative spatial patterns, and b+c is the sum of the voxels uniquely identified in each pattern. A higher κ value denotes a higher degree of overlap, where a score of 0 represents no overlap and a value of 1 represents perfect overlap (Zou et al., 2004).

## 3. Results

### 3.1. Assessing disease progression and motor symptom profile

#### 3.1.1. Disease progression (survival rates) and motor symptom duration

As expected, we observed lower survival rates for Hu-PFF-injected mice, however, we observed that male Hu-PFF-injected mice reached their humane endpoint prior to the end of the experiment (120 dpi). This significant injection group by sex interaction (p = 0.00166; Cohen’s d = −3.25) suggests a significant physiological difference in response to the Hu-PFF injection in males and females (Figure 2A). Male Hu-PFF mice reached their humane endpoint after an average of 23 days of overt motor symptomatology.

**Figure 2.**
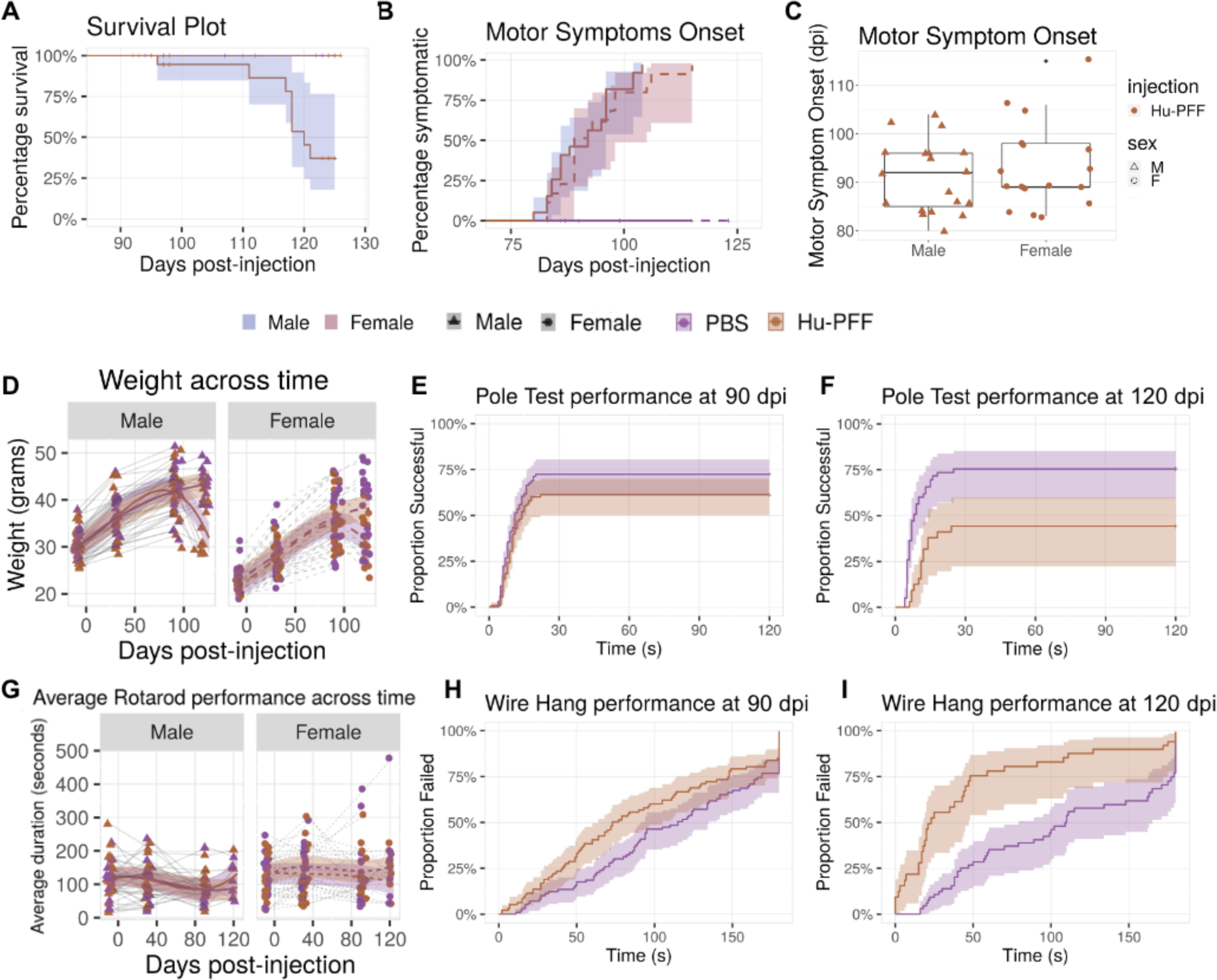
Sex-focused examination of disease progression and motor symptomatology. [A] Survival plot for M83 hemizygous mice. Percent survival of the mice plotted across the days post-injection (dpi) with the experimental endpoint being 130 dpi. Almost 30% of the male Hu-PFF-injected mice succumbed to their symptoms/disease progression a couple days prior to the 120 dpi time point. Group differences in survival rate were examined using general linear models, and a significant group by sex interaction was observed whereby male Hu-PFF-injected mice had lower survival rates (p=0.00166). Of the mice that did not survive within the time frame examined (<130 dpi), the mice sustained motor symptomatology for on average 23 days. [B] Percentage of mice showing motor symptoms across the number of days post-injection. No significant differences between the sexes were observed with regards to the percentage of mice being symptomatic. [C] With regards to motor symptom onset, in terms of the days post-injection, no significant differences were observed between the sexes. [D] Weight trend across disease progression. Significant inverted U-shaped trajectory for Hu-PFF-injected mice, with weight loss as of 90 dpi (compared to PBS-injected mice) (p=0.0056), regardless of sex. [E,F] Pole Test performance at 90 (left) and 120 (right) dpi. Hu-PFF-injected mice had lower proportions of mice successfully passing the test, and took significantly longer to descend the pole compared to their saline injected counterparts (p=0.0304; p=0.017). [G] Average rotarod performance across time showed no significant differences between injection groups and sex. [H,I] Wire-hang performance at 90 (left) and 120 (right) dpi. Significant difference in the proportion of mice that failed (<3 minutes) between injections groups (p=0.0104; p=0.000504), with higher rates of failure for the Hu-PFF-injected mice (red dashed line); no sex difference observed. Purple colour denotes PBS-injected mice and orange colour denotes Hu-PFF injected mice. Line type was used to denote each of the sexes: solid line (with blue shading) for male and dashed line (with red shading) for female mice, except when no sex differences are displayed solid lines then denote both sexes grouped together. Data point shapes also denote the sex of the mice: triangle for males and round for females.

#### 3.1.2. Motor symptom onset

Given the differences in sex-specific survival rate, we assessed differences in the length of time required for the mice to be considered symptomatic; however no sex-specific differences were observed (Figure 2B). Similarly, no significant sex differences in the number of days after which motor symptoms were first observed between the two injection groups were observed. (Figure 2C). Contrary to our survival analysis, our findings do not suggest a sex difference in the onset and rate of onset of overt motor symptomatology.

#### 3.1.3. Motor symptomatology

Beyond observational motor symptoms, we examined motor behaviour and disease progression via weight tracking and motor performance. For the M83 mouse model, initial phenotypic changes include neglect of grooming, weight loss, and reduced ambulation (Giasson et al., 2002). We observed a significant inverted U-shaped trajectory for Hu-PFF-injected mice, compared to the consistent weight increase in PBS-injected mice (p=0.00108; d=-3.31). In the M83 mice injected with Hu-PFF, weight peaks at ∼90 dpi then declines steadily thereafter. However, no sex-specfic differences were observed when examining the triple interaction between group, sex, and time since injection (Figure 2D).

At each timepoint, performance on the rotarod, wire hang and pole test were assessed. Average rotarod performance across time showed no significant differences for the sex by injection group by time interaction (Figure 2G); nor were there group by sex differences in performance at 90 and 120 dpi. Performance was evaluated cross-sectionally at each timepoint for both wire hang and pole tests in terms of failure/success rates, and performance duration. Hu-PFF-injected mice had significantly worse performance (higher failure rates and shorter hang duration) at both 90 dpi (p=0.0104; hazard ratio [HR] = 1.70) and 120 dpi (p=0.000504; HR=3.665) on the wire hang test compared to PBS (Figure 2E-F). Similarly, for the pole test, these mice had significantly worse performance (lower proportion of successfully passing the task and longer time to descend the pole) at 90 dpi (p=0.0304; HR=0.603) and 120 dpi (p=0.017; HR=0.229) (Figure 2H-I).

### 3.2. Longitudinal volumetric analysis

We observed significant differences in the rate of volume change as time progressed for Hu-PFF in comparison to PBS mice (Figure 3A). Similar to our previous findings, these significant differences were mainly observed in the key structures implicated in synucleinopathies (Tullo et al. 2023), as well as with known connections to and from the injection site (right caudoputamen); namely, in voxels in the contralateral striatum, bilateral somatomotor cortices, periaqueductal gray, as well as highly connected regions such as voxels within the bilateral thalami, all surviving 1% False Discovery Rate (FDR) (Benjamini & Hochberg, 2000).

**Figure 3.**
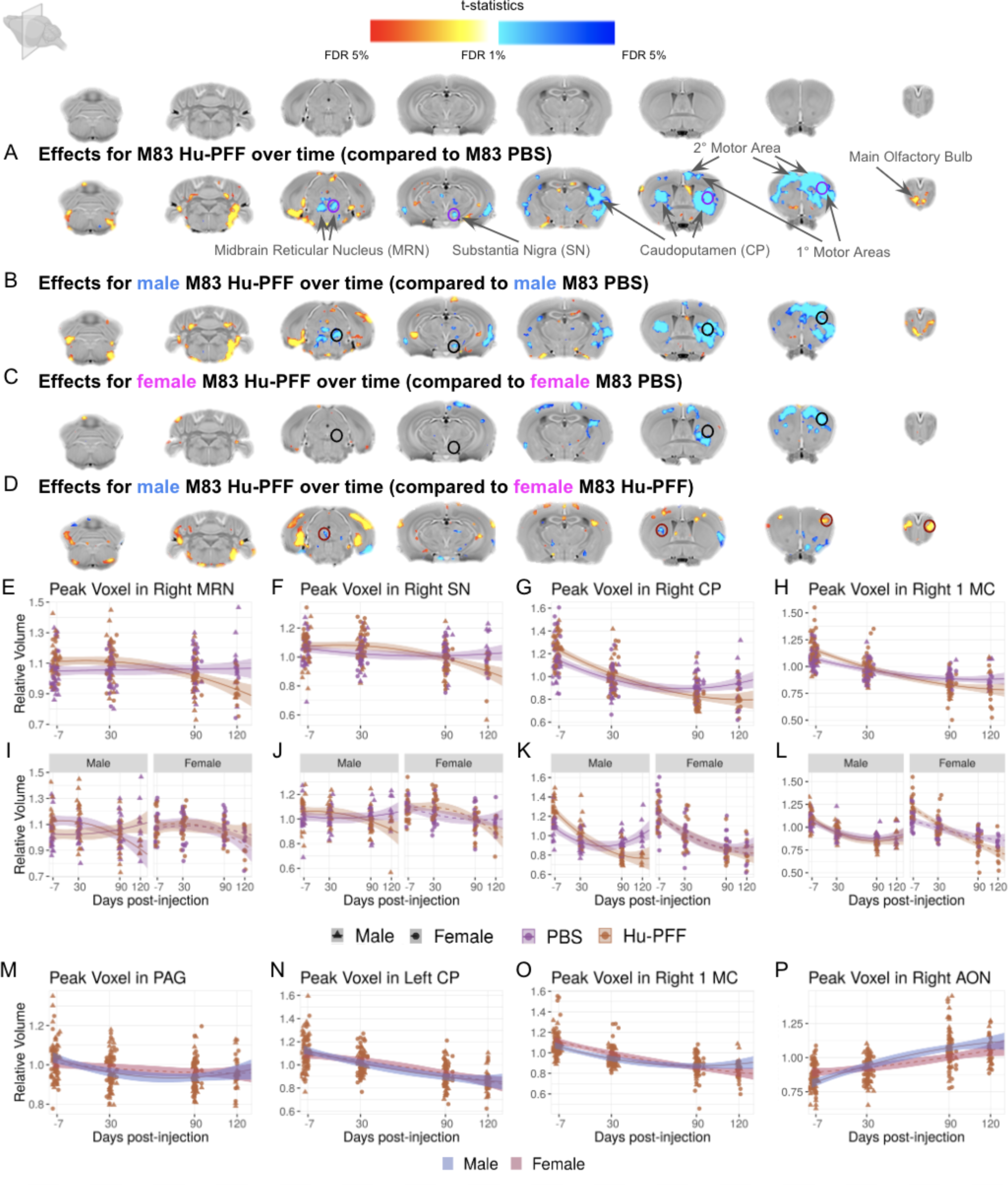
Sex differences in Hu-PFF-induced brain atrophy examined using voxel-wise volumetric trajectories over time. [A-D] Coronal slices of the mouse brain (from posterior to anterior) with t-statistical map overlay; demonstrating [A] the effects of injection over time in M83 hemizygous mice, [B] the effects of injection over time in male mice, [C] the effects of injection over time in female mice, [D] the effect of sex over time for Hu-PFF-injected mice. Colour map describes the direction of the t-statistics; cooler colours denoting negative values; most commonly corresponding to volume decline and warmer colours denoting positive values, corresponding to volume increases over time for the group of interest indicated. [E-P] Plot of relative volume change (mm^3^) over the four time points (−7, 30, 90 and 120 days post-injection) for a peak voxel in [E,I] the right midbrain reticular nucleus (MRN), [F,J] right substantia nigra (SN), [F] the injection site (right caudoputamen (CP)), [H,L] right primary motor area (1 MC). Plots E-H describe group differences in volume over time, while plots I-L describe such differences with respect to each sex. Purple line for PBS-injected mice, orange line for Hu-PFF-injected mice, solid line and triangle points for male and dashed line and circular points for female mice. Plots M-P describe sex differences in volume over time for the Hu-PFF-injected mice; orange points and line describe these Hu-PFF injected mice, where solid line, triangle points, and blue shading describes the male while the dashed line, circular points and red shading describe the trajectory for the female mice. Overall, volumetric decline was observed for Hu-PFF injected mice (compared to PBS-injected mice), with steepest rate of decline for male Hu-PFF-injected mice.

#### 3.2.1. Sex-specific volumetric trajectories

There were significant sex-specific differences in volumetric change over time. We observed a significant three-way interaction of the injection group by sex by time, with the fastest rates of neurodegeneration occurring in male Hu-PFF-injected mice. This accelerated rate for the males was localized in voxels within the injection site (right striatum) (Figure 3G,K), the ipsilateral substantia nigra (Figure 3F,J), bilateral motor cortices (primary and secondary motor areas) (Figure 3H,L), and midbrain areas such as the midbrain reticular nucleus (Figure 3E,I). We explicitly examined injection group differences over time for each sex separately to better assess sex-specific influence of Hu-PFF injections (Figure 3B-D). Qualitatively, we observed greater differences between Hu-PFF- and PBS-injected mice for the male mice compared to the same comparison in female mice (Figure 3B-C). Such differences between each sex-specific comparison can be best seen in the caudoputamen both ipsi- and contralateral to the injection, and is particularly striking for more posterior areas such as midbrain and brainstem areas. Accordingly, we examined sex differences for the Hu-PFF-injected mice over time (Figure 3D), and we observed steeper rates of decline for male mice, particularly in regions such as the contralateral caudoputamen (Figure 3N).

### 3.3. Whole brain structural covariance patterns of atrophy

Although our investigation of volumetric trajectories demonstrated significant volumetric declines as a result of widespread toxicity of fibrillar aSyn resulting from Hu-PFF injection, we further sought to investigate patterns of voxels that support a network-like atrophy pattern as a result of the prion-like spreading of aSyn.

#### 3.3.1. Peri-motor symptom onset

We retained k=6 components for analysis of covariance patterns at 90 dpi (as determined by stability analysis; see section *7.8. Orthogonal projective non-negative matrix factorization* in Supplementary Materials; Supplementary Figure S2). Although our general linear models for each of the 6 components revealed no significance with sex by injection group interaction when examining the subject weights (Figure 4A), only one of the six components statistically separates the Hu-PFF-from PBS-injected groups (component 1; p=0.0323; d=2.179). The spatial covariance pattern consisted of voxels in regions with known connections to the injection site as demonstrated by the striatal-pallidal-midbrain patterning, with strong cortical and thalamic involvement (Figure 4A). All 6 components are detailed in Supplementary Figure S3. Furthermore, for this component (component 1), we also observed a significant effect of sex (p=2.18e-8; d=6.23), with higher subject weights for all females (Figure 4B). Given that all mice are on an M83 background, these findings provide support for further examining sex-specific structural covariance patterns.

**Figure 4.**
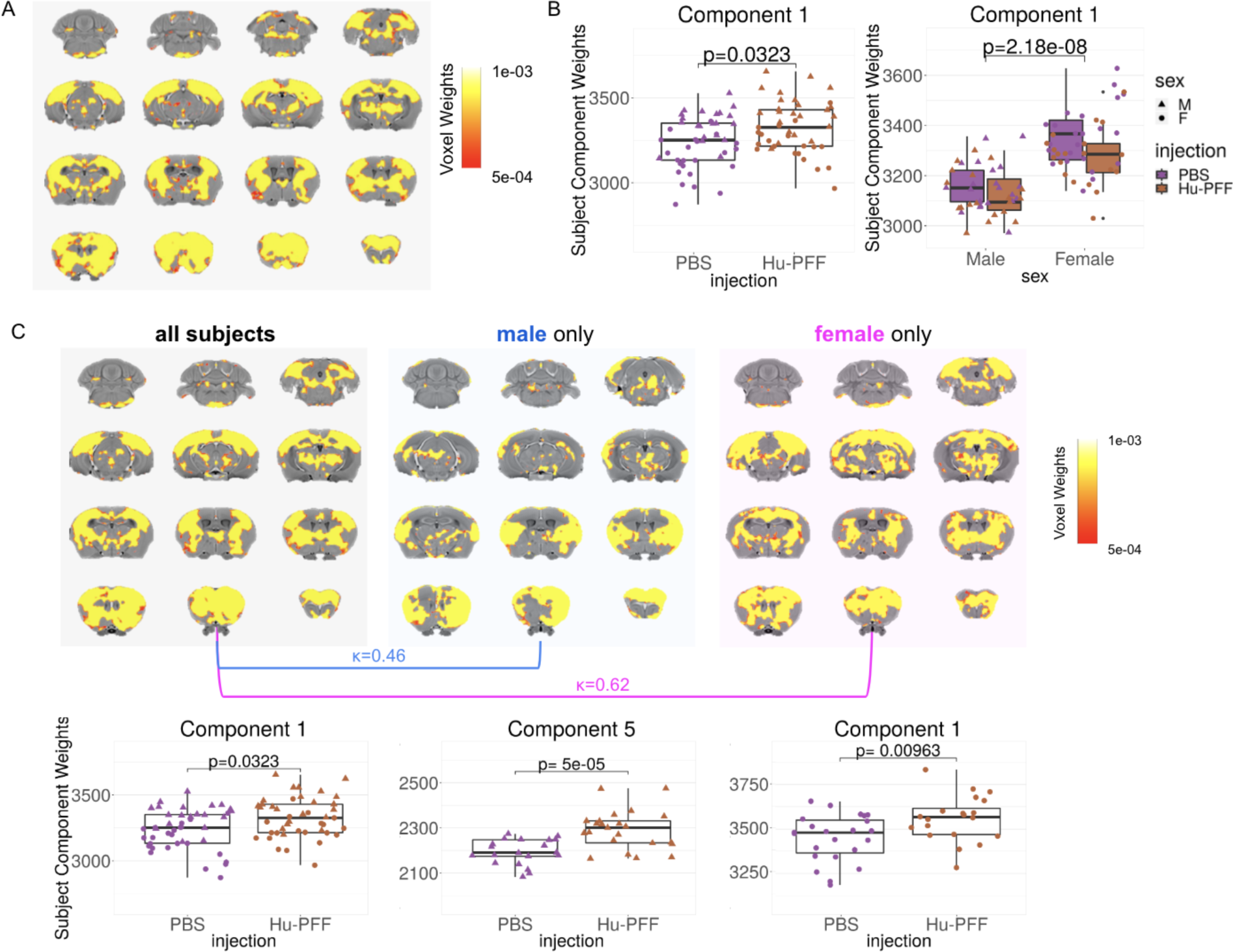
Sex driven patterns of pathology at peri-motor symptom onset. [A,C] The spatial pattern of voxel component weights (denoted by the colour map) plotted onto the mouse brain, depicting patterns of voxels sharing a similar variance pattern. [A-C] NMF decomposition at 90 dpi (peri-symptom onset) for all subjects (n=87). [B] Plot of subject weights shows significant main effect of injection group (p=0.0323) and main effect of sex (p=2.18e-8), but no significant interaction. Higher subject weights for Hu-PFF-injected mice (compared to PBS-injected nice) and for females (compared to males) for component 1 weights. [C] Sex specific NMF runs males only (n=44) (middle) versus females only (n=43) (right); k=6 components, comparison versus all subject NMF run (left; shown as well in [A]). Significance for aSyn Hu-PFF-induced network level pathology assessed using general linear models; significant group differences in subject component weights were observed for only one of six components for each of the three NMF runs (all subjects: component 1; male only: component 5; and female only: component 1). Dice-Kappa overlay score was used to examine the spatial similarity between the three spatial patterns, such that a higher Kappa denotes greater overlap. The greatest overlap with the all subject NMF component is between it and the female-only spatial component (κ=0.62), thus suggesting that the all subject component is predominantly female driven, despite observing no statistical group by sex interaction significance of the subject weights for the all subject component 1. Purple for PBS-injected mice, orange for Hu-PFF-injected mice, triangle points for male and circular points for female mice.

For each sex-specific covariance pattern, we similarly observed only one of the six components with significantly higher subject weights for the Hu-PFF-injected mice, compared to their saline counterparts (component 5 for the male only analysis (p=5e-5; d=0.2445) and component 1 for the female only (p=0.00963; d=2.716) (see Supplementary Figure S4 and S5 for all 6 components in each sex-specific analysis).

Both sex-specific Hu-PFF-induced patterns consisted of voxels within the same regions; this included: voxels in the basal ganglia and thalamus, anterior to posterior dorsal cortical areas, and in the midbrain and pons (Figure 4C) and were consistent with the all subject OPNMF pattern described above. Dice-kappa overlap scores revealed a high degree of overlap between the females and the all subjects pattern (κ=0.62) comparatively to the male only overlap (κ=0.46) (Figure 4C). These findings suggest that, in contrast with our univariate analysis that reveals more aggressive localized neurodegeneration in male mice, there is a more spatially widespread neurodegeneration pattern for the female mice.

#### 3.3.2. Post-motor symptom onset

Analysis of the 120 dpi (once overt motor symptoms are present) yielded components 3 and 6 which differentiated the Hu-PFF from PBS injected groups. Unfortunately, we are underpowered to explicitly examine sex-specific covariance patterns given the high numbers of attrition due to disease progression (with <8 Hu-PFF male mice remaining). Nonetheless, we did observe a trend-level injection group by sex interaction (p=0.088; d=1.764) for component 6 only, with slightly higher subject weights for male Hu-PFF-injected mice (Supplementary Figure S6). More information can be seen in the Supplementary material, section *7.10. Post-motor symptom onset whole brain structural covariance patterns of atrophy* and Supplementary Figure S6).

### 3.4. Western Blot

We next sought to examine whether there were inherent sex differences in the M83 model in the expression of the human aSyn transgene, as well as the normally expressed mouse aSyn that may represent a baseline difference rather than a difference in pathological accumulation. The levels of aSyn serine129 phosphorylation (pS129), a classical pathological modification of aSyn (Altay et al., 2023; Anderson et al., 2006; Fujiwara et al., 2002; Lashuel et al., 2022), did not differ between males and females when comparing three different brain regions (striatum, brainstem, and cortex) (Supplementary Figure S7). Similarly, human and total (human + mouse) aSyn did not differ between the sexes (Supplementary Figure S7).

Additionally, we analyzed the RIPA-insoluble fraction of these same samples, since aSyn aggregates can be found in this fraction. The signal obtained for pS129 was weak (Supplementary Figure S7), which is expected for healthy mice (Elfarrash et al., 2019; Lackie et al., 2022), and we found no sex differences for all variants of aSyn (pS129, human, and mouse) (Supplementary Figure S7).

## 4. Discussion

The findings of this study provide valuable insights into the complex interplay between biological sex, disease progression, and neuroanatomical alterations in synucleinopathies. Here, our investigations encompassed multiple aspects of disease pathogenesis and aSyn spreading over time, including longitudinal neuroanatomical alterations, structural covariance patterns of atrophy, and motor symptomatology assessment.

Neuropathological examination of the brain at different stages of PD progression has been extensively documented, and the revealed patterns of pathology heavily suggest aSyn spreading between anatomically connected regions over time in what have become classic studies of this phenotype (Braak et al., 2003). Despite extensive examination of aSyn spreading in animal models of synucleinopathy (Bétemps et al. 2014; Chu et al., 2019; Desplats et al., 2009; Fares et al., 2016; Froula et al. 2019; Luk et al., 2012a; 2012b; Karampetsou et al., 2017; Masuda-Suzukake et al., 2013; 2014; Mougenot et al., 2012; Panattoni et al., 2018; Paumier et al., 2015; Sacino et al. 2014; Tapias et al., 2017; Watts et al., 2013) via widespread Lewy body-like deposition in the brain, affirmative evidence of atrophy being caused by this hypothesised pathogenic process is still lacking. Here, we examined longitudinal within subject measures of pathology over the disease time course in the same model of synucleinopathy from pathology seeded at a known central locus across the whole brain. Our findings agree with our previous cross-sectional study (Tullo et al., 2023) that demonstrated that aSyn mediated neurotoxicity preferentially impacted regions highly interconnected with the injection site displayed the most severe pathology, such as basal ganglia areas and connected motor cortices. Most importantly however, in this study we are better suited to examine spreading and progressive pathology, and assess normative variations in disease progression, both in terms of brain pathology and motor symptomatology. Our results highlight the significance of sex-specific differences in the context of aSyn spreading and neurodegenerative disease modeling by presenting sex-specific spatial and temporal aspects of synucleinopathy-associated disease progression.

We observed more aggressive neurodegeneration (atrophy) in male Hu-PFF-injected mice. Moreover, the identification of sex-specific patterns of structural covariance at both peri-motor symptom onset and post-motor symptom onset time points underscores the need for a nuanced approach when studying disease progression and may suggest multiple sex-specific mechanisms at play. While both male and female mice displayed Hu-PFF-induced patterns of atrophy, we observed early widespread pathology in the female mice that seems to be less toxic given that these mice survived longer than their male counterparts. However, once the mice showed overt motor symptomatology, the male Hu-PFF-injected mice displayed widespread pathology which coincided with their lower rates of survival (with higher rates of male Hu-PFF-injected mice reaching their humane endpoint prior to the experimental endpoint). The levels of host aSyn (mouse or human) and baseline phosphorylated S129 aSyn (the substrate for aSyn spreading and toxicity) were not different between sexes. Hence, it is likely that intrinsic sex-specific molecular, cellular or circuit differences in aSyn spreading and toxicity contribute to the observed differences in brain atrophy and survival. Whether these are hormonally regulated remains to be defined. This study represents a critical step forward in understanding the impact of sex on disease progression in preclinical M83 Hu-PFF mouse models of synucleinopathies. Our findings emphasize the importance of considering sex-specific pathologies for investigating disease mechanisms and therapeutic interventions.

Our findings not only underscore the importance of considering sex (and gender for human studies) differences in synucleinopathies but also reveal a gap in the existing literature. The scarcity of studies that systematically focus on such differences in these disorders is evident, despite reported and anecdotal phenotypic differences between men and women diagnosed with synucleinopathies. Overall, men are more likely than women to experience a higher prevalence, increased incidence, greater disease severity, and heightened susceptibility to synucleinopathies (Coon et al., 2019; Picillo et al., 2017; Utsumi et al., 2020; Yamamoto et al., 2014; Zhou et al., 2015). In terms of neuroanatomical changes between the sexes, unfortunately there is a lack of neuroimaging studies centered on sex differences in synucleinopathies and in the vast PD literature in general, in spite of the overwhelming clinical and epidemiological evidence supporting sex differences in diagnosis, presentation, and prevalence (Salminen et al., 2021). Namely, three structural MRI studies have explored sex differences in gray matter brain atrophy within the context of PD. In studies of cortical thickness, one study by Tremblay et al. (2020) in *de novo* patients with PD did not observe any sex differences, whereas work by Oltra et al. (2022) observed significant cortical thinning in various regions across the brain (including all four lobes: frontal, parietal, temporal, and occipital lobes) in male compared to female patients, similar to work by Yadav et al. (2016) in patients undergoing treatment for PD. With regards to whole brain volume measures, using DBM, all three studies reported greater male atrophy in patients. Specifically, they observed more cortical regions with male-driven (eleven) vs female-driven atrophy (six), and greater subcortical atrophy in males (Yadav et al., 2016; Tremblay et al., 2020; Oltra et al., 2022). Nonetheless, more evidence is needed to effectively parse through sex-specific differences in neuroanatomical pathology to be able to use neuroimaging techniques as valuable biomarkers for stratifying patients based on their risk, rate and severity of disease progression (Meoni et al., 2020).

The predominant notion driving these sex differences is thought to be the role of hormones and sex chromosomes (Martinez-Martin et al., 2012; Silva et al., 2008; Yamamoto et al., 2014). However, there is a lack of evidence for the associations between events of hormonal fluctuations in women (such as menstruation onset age, menopause onset age, fertile lifespan, pregnancy history, use of oral contraceptives and hormone replacement therapy) and the risk of developing PD in women (Brakedal et al., 2022; Castelnuovo et al., 2020; de Lau et al., 2014). Thus, there is still a lack of causality for the neuroprotective properties of estrogen (Costantino et al., 2022). In fact, as stated in a recent systematic review on sex differences in movement disorders by Raheel et al. (2023), several important questions remain unanswered with regards to the estrogen conversation: disentangling endogenous versus exogenous estrogen exposure, the causality of estrogen effects on different aspects of the disease, and the intricate interplay between hormonal changes and the progression of synucleinopathies, determining the threshold, time window (e.g., premenopause versus perimenopause versus postmenopause), and other potential modifying factors that influence the neuroprotective effect of estrogen (Raheel et al., 2023).

Despite the insights provided by this study, it is essential to acknowledge its limitations. First, this study primarily focused on structural changes, and future research should investigate functional and molecular aspects of sex-specific differences in synucleinopathies. Nonetheless, this manuscript provides a much-needed characterization of the neuroanatomical pathological changes over the disease time course in vivo, similarly to human synucleinopathy imaging studies. Beyond this characterization, our specific focus on both sexes and sex differences allows us to examine sex-specific disease presentation and progression otherwise uncharacterized in this well-used model. Our findings advocate strongly for a shift in pre-clinical research practices to consistently include both male and female model systems to better model more than half the world’s population. By acknowledging and addressing sex differences with investigation in both female and male mice, we aimed to enhance the translatability of these models, improving its face and construct validity. Nonetheless, further research is warranted to elucidate the underlying mechanisms driving these sex-specific variations, as the identification of the molecular pathways driving these differences may open new avenues for sex-specific therapeutic strategies, often neglected in many research avenues. Similarly, beyond the importance of using both female and male mice for preclinical testing, given that the age of onset of such neurodegenerative diseases occurs around peri- and post-menopause, it has become clear that our modelling of the female condition in mice should include the hormonal fluctuations occurring during these phases to most accurately disentangle the role of hormones in disease pathogenesis. We thereby urge such investigations to also be considered. This paradigm shift is vital to drive advances in the field, improve our understanding of sex-specific mechanisms in synucleinopathies, effective translation from animal models to human organisms, and to pave the way for the development of novel, sex-specific therapeutic strategies. Ultimately, such efforts will contribute to the establishment of personalized medicine approaches that consider the unique needs of all patients affected by these devastating neurodegenerative diseases.

A final, yet important, limitation is the investigation of overt motor symptomatology when observing the mice in their home cage. These symptomatologies have been extensively characterized in this model over the years (Bétemps et al., 2014; Froula et al., 2019; Luk et al., 2012a; Luk et al., 2012b; Masuda-Suzukake et al., 2013; Masuda-Suzukake et al., 2014; Mougenot et al., 2012; Sacino et al., 2014; Watts et al., 2013). However recent evidence has emerged suggesting cognitive impairment may precede the motor symptomatology as evidenced by touchscreen cognitive tasks performed at pre-motor symptom onset time points (∼50-60 days post-injection) (Lackie et al., 2022; Tullo et al., 2023). This investigation of nonmotor symptomatology is important with regards to the clinical presentations of synucleinopathies wherein nonmotor symptoms (autonomic, olfactory, etc.) present prior to diagnosis (Boeve et al., 2003; Kao et al., 2009; Pagonabarraga & Kulisevsky, 2012; Sveinbjornsdottir, 2016). Future work should examine symptoms onset with regards to both motor and nonmotor symptomatology.

## 5. Conclusions

In conclusion, this study provides a clear example of why it is important to incorporate sex-specific considerations into preclinical mouse models of synucleinopathies. Our findings provide compelling evidence of sex-specific differences in the spatiotemporal dynamics of synucleinopathy-associated pathology and disease progression, underscoring the importance of considering sex-specific factors in research and clinical practice. Future research should aim to investigate the molecular and cellular mechanisms driving sex differences in synucleinopathy pathogenesis, as it may offer significant implications for patient management and the development of targeted therapeutic interventions. Furthermore, further investigation into diverse hormonal models could better capture the nuanced impact of these factors on disease susceptibility and presentation. Embracing this paradigm shift will be essential for advancing our understanding of disease mechanisms and developing tailored therapeutic strategies, bringing us closer to personalized medicine approaches for patients affected by synucleinopathies and other neurodegenerative diseases.

