## Supplementary materials for "Sex-focused analyses of M83 A53T hemizygous mouse model with recombinant human alpha-synuclein preformed fibril injection identifies female resilience to disease progression: A combined magnetic resonance imaging and behavioural study"

*7.1. Animals*

All mice were bred in-house via the following breeding scheme: TgM83+/- x TgM83+/-. Mice were housed at the Douglas Research Center Animal Facility (McGill University, Montreal, QC, Canada) under standard housing conditions with food and water *ad libitum*. Mice were typically housed in maximum groups of four, and mice were rarely housed singly unless fighting occurred (only seen in males). Mice were housed with a 12/12-hour light/dark cycle, with lights on at 08:00.

*7.2. Injection materials and Stereotaxic injections*

The Hu-PFF were made and characterized by the Early Drug Discovery Unit (EDDU) at the Montreal Neurological Institute using established standard operating procedures (SOPs) (production: https://zenodo.org/record/3738335#.Yp46fXbMKUl; characterization: https://zenodo.org/record/3738340#.YzIjBezML0p) (Volpicelli-Daley et al., 2014).

Hu-PFF were sonicated and DLS analysis was completed to ensure the average diameter of PFF was <100 nm. PFF were added to 200 mesh copper carbon grid (3520C-FA, SPI Supplies), fixed with 4% PFA for 1 minute and stained with 2% acetate uranyl (22400-2, EMS) for 1 minute. PFF were characterized using a negative staining protocol (https://zenodo.org/record/3738340#.Yp46xnbMKUl) and visualized using a transmission electron microscope (Tecnai G2 Spirit Twin 120 kV TEM) coupled to an AMT XR80C CCD Camera, and analyzed with Fiji-ImageJ 1.5 and GraphPad Prism 9 software. Electron microscope characterization of the fibrils in terms of distribution per length can be seen in Supplementary Figure S1.

The mice were anaesthetized with isoflurane (5% induction, 2% maintenance), given an injection of carprofen (0.1cc/10 grams) and xylocaine was applied on the scalp for pain relief, and the mice were positioned into a stereotaxic platform. Recombinant human alpha-synuclein fibrils (total volume: 2.5 µL; 5 mg/mL; total protein concentration, 12.5 µg per brain) were stereotaxically injected in the right dorsal striatum (co-ordinates: +0.2 mm to +0.2 mm relative to Bregma, +2.0 mm from midline, +2.6 mm beneath the dura). The control condition was M83 mice receiving sterile PBS in the same injection site. Injections were performed using a 5 µL syringe (Hamilton; 33 gauge) at a rate of 0.25 µL per min with the needle in place for about 5 min prior to and after infusion of the inoculum. Different syringes were used for each type of inoculum to prevent any contamination. Post-injection, the mice were placed on a heating pad for recovery before being returned to their home cage.

After inoculation, the mice were monitored weekly for health and neurological signs such as reduced grooming, kyphosis, and/or decreased motor functioning (including reduced ambulation, tail rigidity, paraparesis). The frequency of monitoring increased upon the onset of the symptomatology up until the experimental or humane endpoint.

*7.3. MRI acquisition: anaesthesia protocol*

For each time point acquisition, the mice were anaesthetized with isoflurane at 3% induction (with 1% oxygen flow rate) for 3:30 minutes followed by a bolus intraperitoneal injection of dexmedetomidine (1:240 dilution). The mice remained in the induction chamber until 5:30 minutes elapsed by which the mice were transferred to the MRI scanner and placed under 1.5% isoflurane with a constant infusion of dexmedetomidine (0.05 mg/kg/h).

*7.4. Pre-processing*

All brain images were exported as DICOM from the scanner and converted to the MINC (Medical Imaging NetCDF) file format. Image processing was performed using the MINC suite of software tools (http://bic-mni.github.io). At this stage, the images were manually inspected for artifacts (hardware, software, motion artifacts or tissue heterogeneity and foreign bodies) and images with said artifacts were excluded (https://github.com/CoBrALab/documentation/wiki/Mouse-QC-Manual-(Structural)). Injection site was verified for accuracy by examining the resulting edema of the surgery present in the brains at the 30 dpi scan time point. Notably, no remaining edema is present for the subsequent scan time points beyond the 30 dpi time point for any of the mice.

After passing quality control, the images were stripped of their native coordinate system, left-right flipped to compensate for Bruker's incorrect DICOM exporting, denoised using patch-based adaptive non-local means algorithm (Coupe et al., 2008), and affinely registered to an average mouse template (the Dorr–Steadman–Ullman atlas; Dorr et al. 2008; Steadman et al. 2014; Ullman et al. 2014) to produce a rough brain mask. Next, a bias field correction was performed, and intensity inhomogeneity was corrected using N4ITK (Tustison et al., 2010) at a minimum spline distance of 5 mm.

*7.5. Deformation-based morphometry*

Given the longitudinal nature of the data, a two-level deformation-based morphometry technique (antsMultivariateTemplateConstruction2.sh; https://github.com/CoBrALab/twolevel_ants_dbm) was used to perform group-wise registration of all MRI data for a single subject to first generate a subject-wise template (first level), followed by the registration of all the subject-wise templates to create a final group-wise template (second level) to enable statistical analysis in a common space. The final deformations fields represent the minimum deformation required, at a voxel-level, to map each subject time point image to the group template. The Jacobian determinants were log-transformed then blurred with Gaussian smoothing using ~0.085 mm full width at half maximum kernel to better conform to normative distribution assumptions for statistical testing (Chung et al., 2001).

Voxel-wise volume measures were derived from two types of Jacobian determinants, depending on the type of analysis. For longitudinal analysis, 'relative' Jacobian determinants (exclusively modelling the nonlinear transformations of the deformations) were used to measure local anatomical differences , whereas 'absolute' Jacobian determinants of the deformations were used to measure all anatomical differences (including the residual global linear transformations attributable to differences in total brain size for example along with the nonlinear transformations) for cross-sectional analyses (specifically, the whole brain structural covariance analyses described in section 2.6.2.1). In both cases, these Jacobians represent expansions (positive values) or contractions (negative values).

*7.6. Behavioural Tasks*

7.6.1. Pole Test

The pole test was administered to examine motor agility and fine motor movements (Matsuura et al., 1997). Each mouse was placed head-upward on the top of a vertical rough-surfaced pole (wooden dowel covered in surgical tape; diameter 8 mm; height 55 cm). The time required for the mouse to descend to the floor was recorded with a maximum duration of 2 minutes. Each subject completed three trials with a 30-minute rest between each trial. We followed the protocol detailed by Matsuura et al. (1997). Using this method, if the mouse descended part way but fell the rest of the way, the behaviour was scored until it reached the floor, however, if the mouse was not able to turn downward and instead dropped from the pole, time was taken as 120 s (maximum value) because of the maximal severity.

7.6.2. Rotarod

To investigate motor coordination and balance, mice were tested on the Rotarod as previously described (Janickova et al., 2017). Mice were placed on the Rotarod (San Diego Instruments; San Diego, CA, USA) and rotation was accelerated linearly from 4 to 25 revolutions per minute (rpm), increasing rpm every 15 seconds. Time spent walking on top of the rod before falling off the rod or hanging on and riding completely around the rod was recorded. Mice were given three trials with a minimum 30-minute inter-trial rest interval.

7.6.3. Wire-Hang

Neuromuscular strength was tested with the wire hang test. The mouse was placed on a wire by waving it gently so that it gripped the wire and then inverted. Latency to fall over the course of three trials was recorded with a 3-minute cut-off time, with a 30-minute inter-trial rest period. A trial of less than 10 seconds was performed again, and trials were concluded at 3 minutes if a mouse was still hanging (Froula et al. 2019).

*7.7. Western Blotting*

Brain tissue from 6-7 months old M83 hemizygous mice was rapidly dissected, flash frozen using dry ice, and stored at -80 °C. Cortical, striatal, and brainstem tissue was homogenized using ice-cold RIPA buffer (50 mM Tris pH 8.0, 150 mM NaCl, 5 mM EDTA,

0.1% SDS, 0.5% Sodium Deoxycholate, 1% Triton-X 100) with phosphatase inhibitors (1 mM NaF and 0.1 mM Na3VO4) and protease inhibitor cocktail (1:100, Catalog#539,134-1SET, Calbiochem). Specifically, a handheld homogenizer with a pestle was used to break down the tissue, followed by three rounds of 7-second sonication at 4 °C, and finally centrifugation for 20 min at 17,000 x g. Supernatant containing RIPA-soluble homogenates was collected, aliquoted, and stored at -80 °C. The remaining pellet was washed with ice-cold PBS, suspended in 4 M Urea and 2% SDS, vortexed for 10 seconds at max speed, sonicated, and centrifuged.

Supernatant concentration was determined using ThermoFisher BCA protein assay kit (Cat# 23227), samples were prepared with 4x sampling buffer (40% Glycerol, 250 mM Tris-HCI, 8% SDS, 4% β-mercaptoethanol, 6 mM Bromophenol blue), boiled at 95 °C for 5 min, and 13ug of protein was loaded into precast 4-12% Bis-Tris gradient gels (ThermoFisher). Protein was transferred onto 0.2 μm PVDF membranes, fixed in 0.4% PFA for 30 minutes at room temperature, stained with 0.1% Amido black, and blocked with either 5% milk or 5% BSA in 1x TBS-T. Membranes were incubated in the following primary antibodies: phospho S129 (1:1000, Cat# ab51253, Abcam, RRID:AB_869973), anti-human a-Syn (1:1000, Cat# ab27766, Abcam, RRID:AB_727020), anti-alpha synuclein (1:1000, Cat# 610787, BD Biosciences, RRID: AB_398108), anti-actin HRP (1:25,000, Cat#A3854, Sigma-Aldrich, RRID:AB_262011). Sheep anti-mouse HRP (1:5000, Cat#SAB3701095, Sigma-Aldrich, RRID: N/A), and goat anti-rabbit HRP (1:10,000, Cat#170–6515, Bio-Rad, RRID: AB_11125142) were used as secondary antibodies. Membranes were exposed using chemiluminescence and imaged the ChemiDoc MP Imaging System. Densitometry was processed and analyzed using Bio-Rad ImageLab software.

#### *7.8. Cox Proportional Hazard modelling*

This cumulative incidence model is considered superior to the t-test or ANOVA given the nature of the data where two measures can be used to denote the behaviour of the mouse. For the survival and motor symptom onset analysis, we examine 1) the average dpi overt motor symptomatology was first observed in ambulating mice or humane endpoint dpi as well as 2) the proportion of the mice at that dpi. For the pole test and wire hang test, Cox modelling is best suited given that these tests are conducted with max latency times (3 minutes success cut-off for wire hang test, and 2-minute failure cut-off for pole test) (Gallino et al., 2019).

For these motor tasks, the Cox Proportional Hazard models tested the effect of the injection groups and sex interaction to explain the rates of successfully completing each motor task (independently) over time, along with trial number and weight as covariates. This model allows for non-parametrically distributed latency times, the unbiased inclusion of failures (trials where mice cannot perform the task due to motor impairment and thus counted as a max time value (for pole test) or obtain a zero value (for wire hang)), and the ability to include covariates and test for their explanatory contributions (Jahn-Eimermacher et al., 2011; Gallino et al., 2019).

#### *7.9. Orthogonal projective non-negative matrix factorization*

OPNMF decomposes an input matrix, dimensions m x n, where m is the number of voxels and n the number of subjects into two matrices. Here, the input matrix was composed of the inverted absolute Jacobian determinants z-scored at each voxel for each subject, loaded as columns. The first output matrix is the component matrix, dimensions m x k, where k represents the number of components (i.e. spatial patterns of covariance). This output matrix is composed of the component scores for each voxel, which describes the groupings of voxels sharing a covariance pattern. With these voxel-wise component weights, the spatial pattern of voxel weights for each component can be plotted onto the mouse brain template generated by DBM. Moreover, with the orthogonality constraint of OPNMF, this variant of NMF allows for the output spatial components to be non-overlapping, and that each output component represents a distinct pattern of voxels. Thus, a specific voxel can only be part of one component, however the voxels within a region can be included in more than one component. The second output matrix is the weight matrix, dimensions k x n, which contains the weightings of each subject for each component, thereby describing how each subject loads onto each pattern. The component and weight matrices are constructed such that their multiplication reconstructs the input data as best as possible by minimising the reconstruction error between the original input and the reconstructed input. General linear models were performed to examine the sex by injection group interaction in the subject weights for each component.

To select the optimal number of components (k) to analyse, we emulated the stability analyses performed in Patel et al. (2020) performed at various granularities (from k=2 to k=20; at intervals of 2). Two measures are commonly examined when assessing stability and determining k. First, accuracy is measured by observing reconstruction error of a decomposition, defined as the Frobenius norm of the element-wise difference between the original and reconstructed inputs. We plot the gradient in reconstruction error, enabling quantification of the gain in accuracy provided by increasing the number of components from one granularity to the next. Second, stability of a decomposition is measured by assessing the similarity of output spatial components across varying splits of subjects. To track stability, OPNMF is performed on various subsets of subjects and the spatial similarity of the resulting outputs describes how consistent results are across the sample. Here, 5 splits were performed at each granularity.

A high stability value with a low gradient in reconstruction error (i.e. a smaller gain in accuracy by increasing the granularity) is optimal. Here, we define optimal as a balance between high stability (indicating the spatial patterns are consistent across subjects), low reconstruction gradient (indicating that the increase in k/added complexity does not confer a large increase in reconstruction accuracy), and low *k* (fewer components represents a more compact deconstruction and less chance of overfitting)  Notably, the gradient in reconstruction error increases as the granularity increases, however at higher granularities there is a greater propensity of overfitting. See Supplementary Figure S2 for plots of stability and gradient in reconstruction error for each OPNMF.

Here, k=6 was selected for the OPNMF performed at both time points. k was selected based on the stability of each decomposition (i.e. the similarity of output spatial components across varying splits of subjects) and the accuracy as measured by the gradient in reconstruction error (Frobenius norm of the element-wise difference between the original and reconstructed inputs), as detailed in Patel et al, (2020).

### Supplementary Results

*7.10. Post-motor symptom onset whole brain structural covariance patterns of atrophy*

In our analysis of the neurodegeneration patterns once overt motor symptoms are visible, at 120 dpi, we observed that components 3 and 6 differentiated the Hu-PFF and PBS injected groups (see Supplementary Figure S6 for all 6 components retained). Component 3 consists of largely the entire bilateral thalamic nuclei, cerebellum, and brainstem regions, as well as some hypothalamic areas while component 6 consists of mainly subcortical regions such as the striatum, pallidum, lateral areas of the hippocampus, as well as midbrain and the lateral ventricles. Given the location injection site, component 6 best depicts a pattern of Hu-PFF-induced spreading as a direct result of the injection. Most importantly however, for component 6, we also observe a trend-level marginally significant injection group by sex interaction ($p=0.088$), where we observe a trend level effect of higher subject weights for male Hu-PFF-injected mice (Supplementary Figure S6). Although sex-specific covariance patterns were generated for the 90 dpi time point, given the high numbers of attrition due to disease progression (with <8 Hu-PFF male mice remaining), there weren't enough mice to perform such analysis at this 120 dpi time point.

### Supplementary Tables

| **-7 & 30 dpi** | | | | | **90 dpi** | | | | | **120 dpi** | | | |
|---|---|---|---|---|---|---|---|---|---|---|---|---|---|
| | **PBS** | **Hu-PFF** | **Total** | | | **PBS** | **Hu-PFF** | **Total** | | | **PBS** | **Hu-PFF** | **Total** |
| **M** | 30 | 34 | **64** | | **M** | 21 | 23 | **44** | | **M** | 9 | 5 | **14** |
| **F** | 31 | 33 | **64** | | **F** | 23 | 20 | **43** | | **F** | 9 | 8 | **17** |
| | | | **128** | | | | | **87** | | | | | **31** |

**Supplementary Table S1. Number of mice per time point for MRI and behavioural testing.** Phosphate buffered saline (PBS); human aSyn preformed fibrils (Hu-PFF); male (M); female (F).

Supplementary Figures

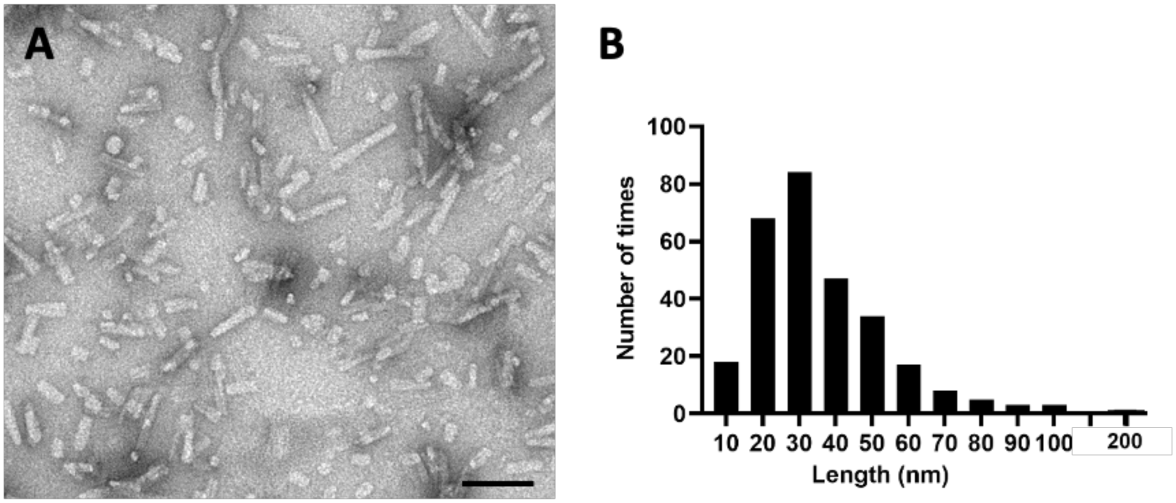


**Supplementary Figure S1. Human alpha-synuclein- preformed fibrils (Hu-PFF) characterization.** [A] Representative photomicrographs of Hu-PFF staining by negative staining and visualized using Tecnai G2 Spirit electron microscope. [B] Histograms showed the Hu-PFF length distribution measured using ImageJ software and their distribution plotted using GraphPad Prism software. Human syn-PFFs sonicated for 30 seconds (n= 288, length average= 35.76nm, median length= 31.65nm, minimal length = 9nm, maximal length= 200nm).

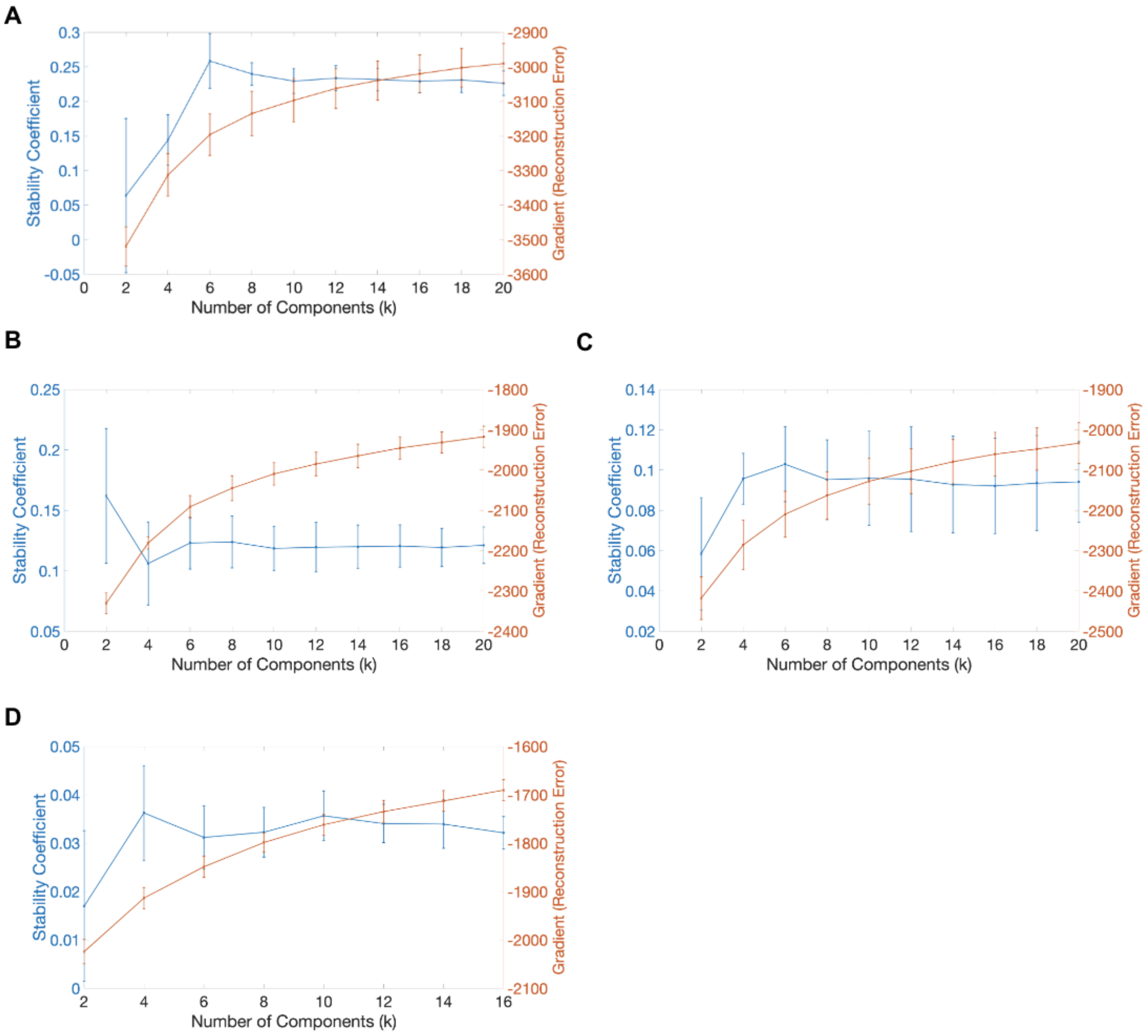


**Supplementary Figure S2. Stability analysis for k=3 to 10 OPNMF component runs.** [A-C] Accuracy and gradient reconstruction error measures at 90 dpi for [A] all subjects, [B] males and [C] female mice respectively. [D] Accuracy and gradient reconstruction error measures for 120 dpi OPNMF run. Gradient in reconstruction error is the quantification of the gain in accuracy provided by increasing the number of components from one granularity to the next. The stability of a decomposition is measured by assessing the similarity of output spatial components across varying splits of subjects; 5 splits were performed at each granularity. For all 4 OPNMF runs, k=6 components were chosen based on the criteria of choosing the highest stability measure with the biggest gain in accuracy (reconstruction error).

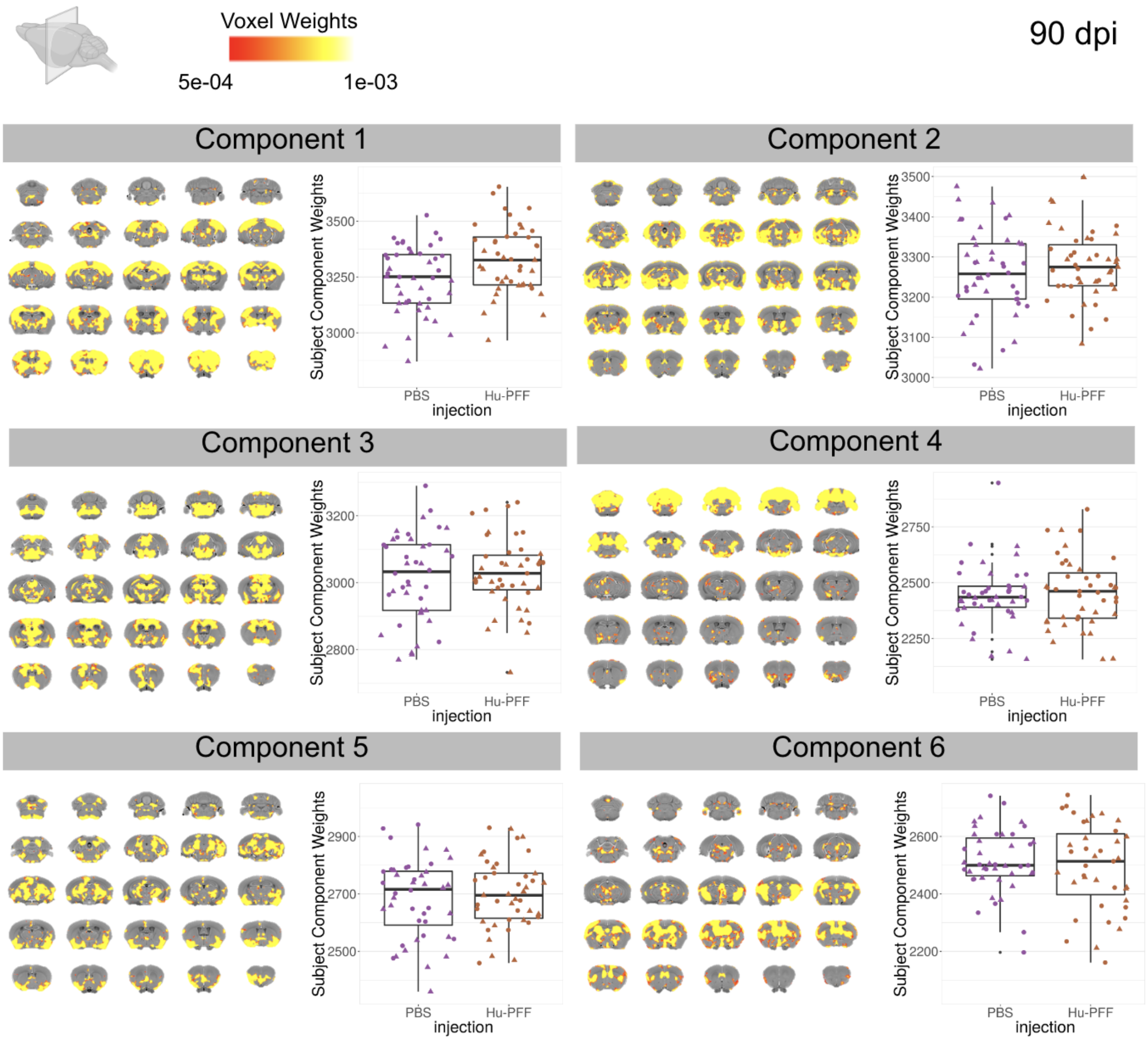


**Supplementary Figure S3. Results of the 90 dpi OPNMF run for all 6 components.** Coronal slices of a mouse brain average displayed from posterior to anterior slices. Colourmap denotes voxel-wise component weights. For each component, the spatial pattern of voxel component scores plotted onto the average mouse brain, depicting the networks of voxels sharing a similar variance pattern (left) and group differences of subject component weightings, describing how each subject loads onto the identified atrophy pattern were assessed using general linear models (right) for each of the 6 components. Component 1 was the only component where the injection group was significantly associated with OPNMF voxel weights. Purple for PBS-injected mice, orange for Hu-PFF-injected mice, triangle points for male and circular points for female mice.

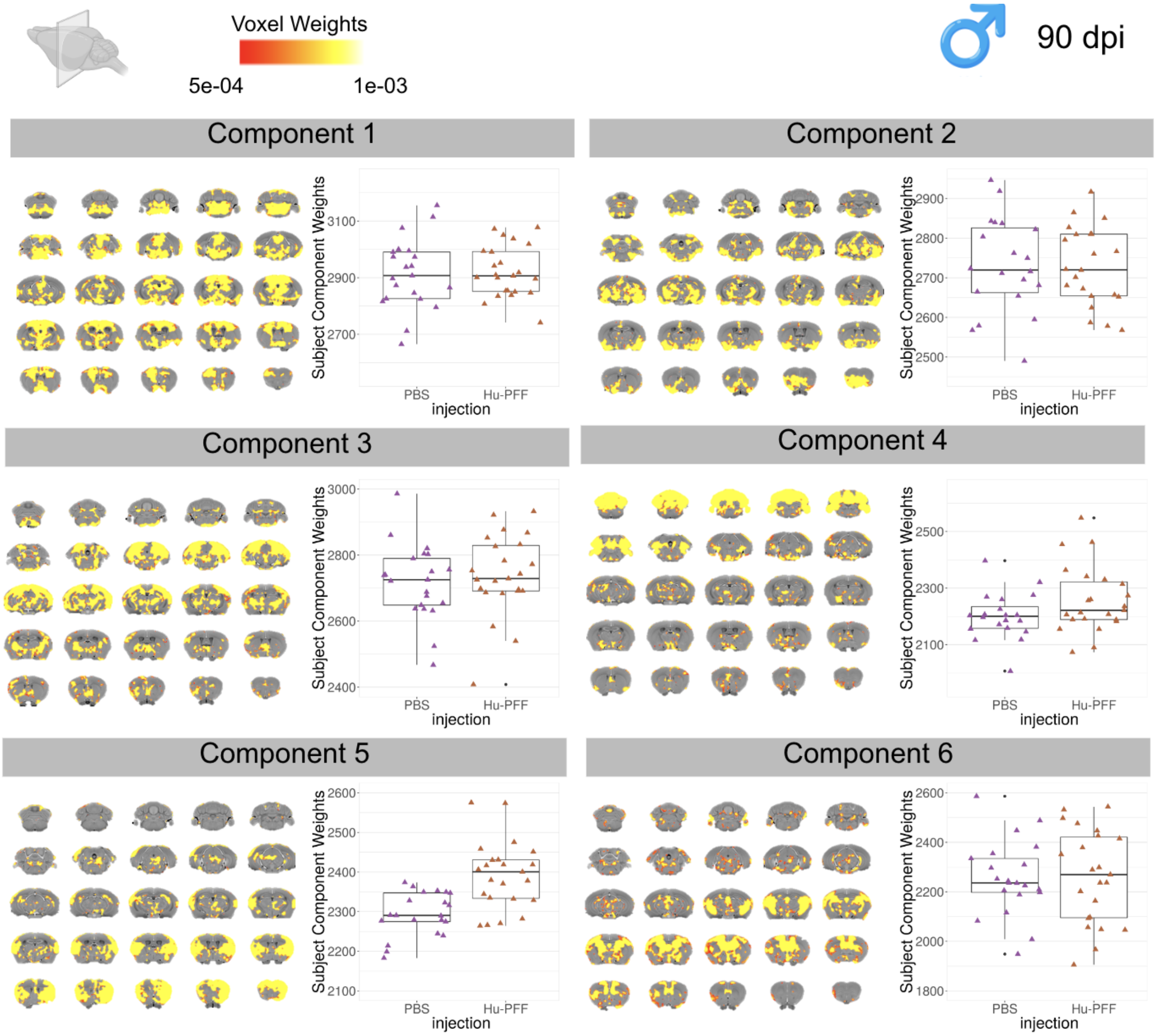


**Supplementary Figure S4. Results of the 90 dpi OPNMF run for all 6 components for M83 Hu-PFF- and PBS-injected male mice.** Coronal slices of a mouse brain average displayed from posterior to anterior slices. Colourmap denotes voxel-wise component weights. For each component, the spatial pattern of voxel component scores plotted onto the average mouse brain, depicting the networks of voxels sharing a similar variance pattern (left) and group differences of subject component weightings, describing how each subject loads onto the identified atrophy pattern were assessed using general linear models (right) for each of the 6 components. Component 5 was the only component where the injection group was significantly associated with OPNMF voxel weights. Purple for PBS-injected mice, orange for Hu-PFF-injected mice, triangle points for male and circular points for female mice.

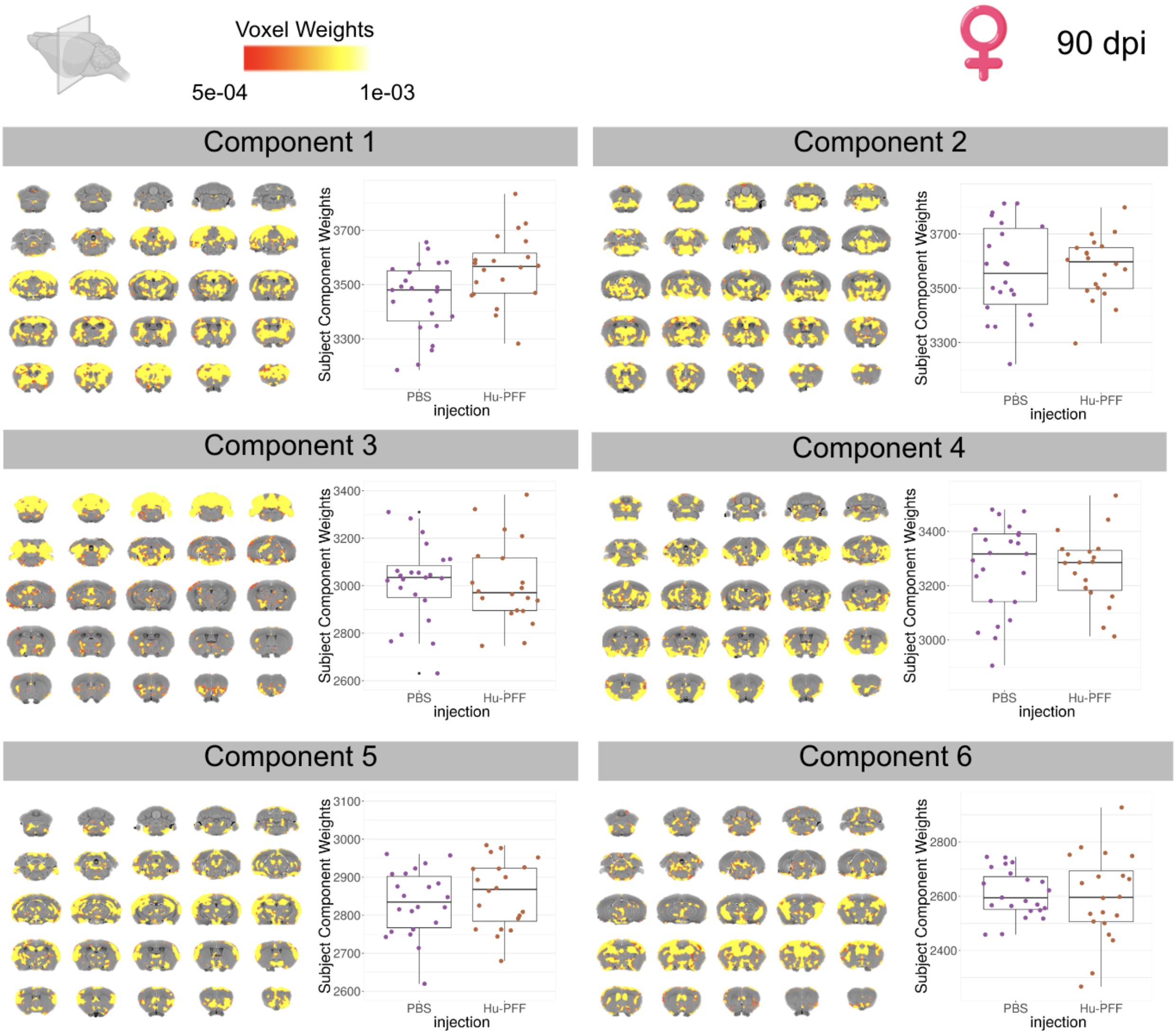


**Supplementary Figure S5. Results of the 90 dpi OPNMF run for all 6 components for M83 Hu-PFF- and PBS-injected female mice.** Coronal slices of a mouse brain average displayed from posterior to anterior slices. Colourmap denotes voxel-wise component weights. For each component, the spatial pattern of voxel component scores plotted onto the average mouse brain, depicting the networks of voxels sharing a similar variance pattern (left) and group differences of subject component weightings, describing how each subject loads onto the identified atrophy pattern were assessed using general linear models (right) for each of the 6 components. Component 1 was the only component where the injection group was significantly associated with OPNMF voxel weights. Purple for PBS-injected mice, orange for Hu-PFF-injected mice, triangle points for male and circular points for female mice.

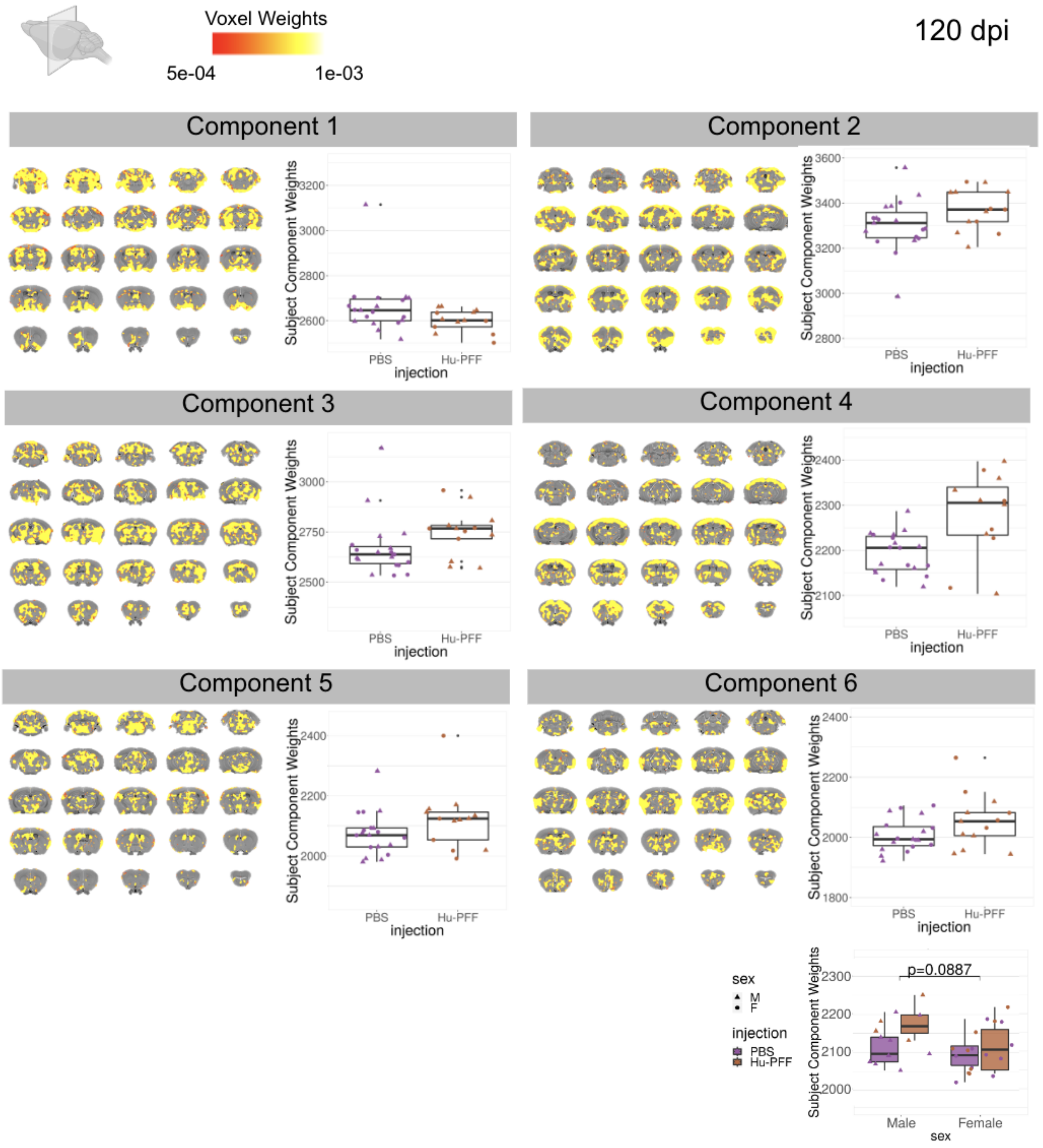


**Supplementary Figure S6. Results of the 120 dpi OPNMF run for all 6 components.** Coronal slices of a mouse brain average displayed from posterior to anterior slices. Colourmap denotes voxel-wise component weights. For each component, the spatial pattern of voxel component scores plotted onto the average mouse brain, depicting the networks of voxels sharing a similar variance pattern (left) and group differences of subject component weightings, describing how each subject loads onto the identified atrophy pattern were assessed using general linear models (right) for each of the 6 components. For components 3 and 6, the injection group was significantly associated with OPNMF voxel weights. Plot of subject weights for component 6 shows marginal significance of injection and sex (p=0.0887), with higher subject weights for male Hu-PFF-injected mice (burgundy triangular markers). Purple for PBS-injected mice, orange for Hu-PFF-injected mice, triangle points for male and circular points for female mice.

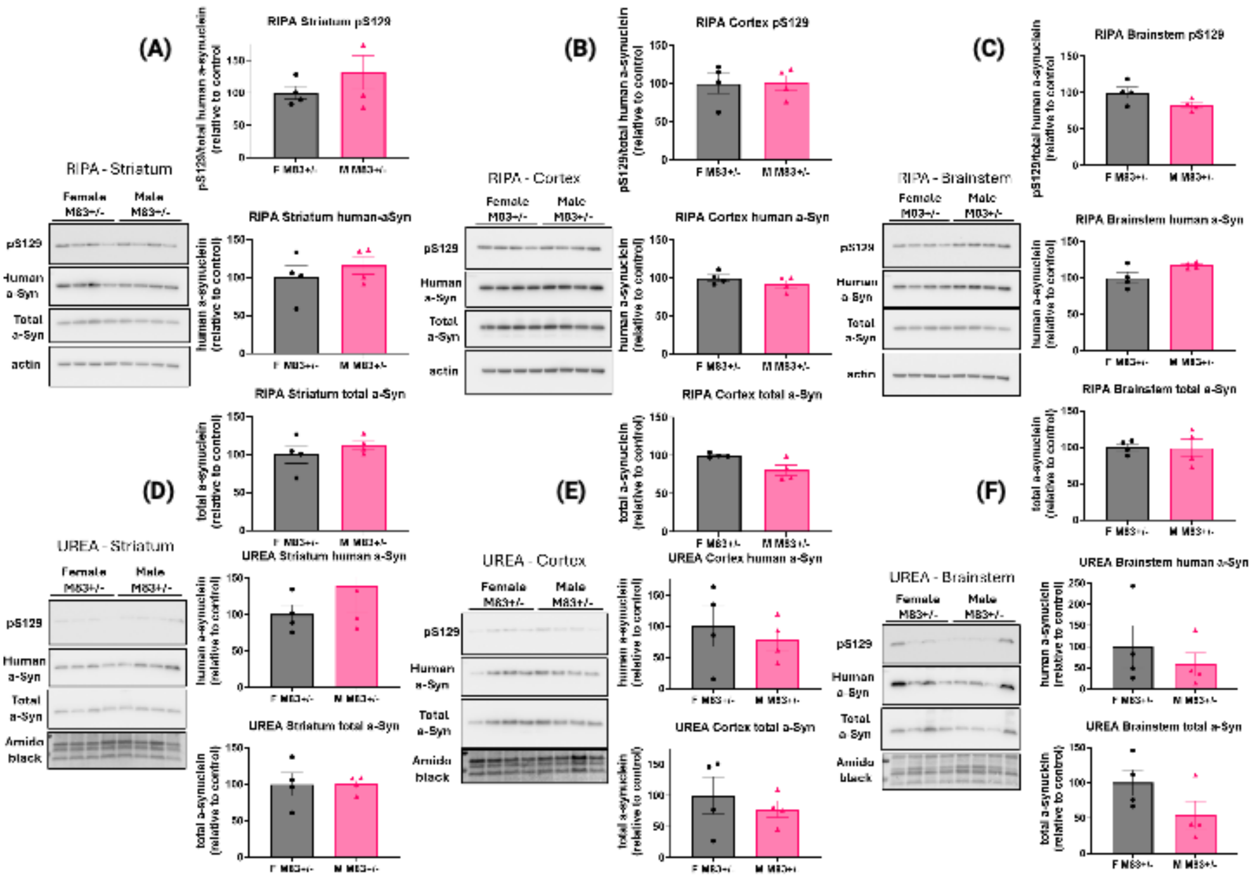


**Supplementary Figure S7. Results of the Western Blotting examining sex-differences.** (A-C) RIPA fraction results for striatum, cortex, and brainstem tissues, respectively. Human, mouse, and phosho129 a-synuclein levels were quantified and analyzed using unpaired t-tests. (D-F) Insoluble fractions obtained (UREA) were run and quantified similarly. No significant differences were observed for either fractions and signals.
